# Characterization of single gene copy number variants in schizophrenia

**DOI:** 10.1101/550863

**Authors:** Jin P. Szatkiewicz, Menachem Fromer, Randal J. Nonneman, NaEshia Ancalade, Jessica S. Johnson, Eli A. Stahl, Elliott Rees, Sarah Bergen, Christina Hultman, George Kirov, Michael O’Donovan, Michael Owen, Peter Holmans, Pamela Sklar, Patrick F. Sullivan, Shaun M. Purcell, James J. Crowley, Douglas M. Ruderfer

## Abstract

Genetic studies of schizophrenia (SCZ) have now implicated numerous genomic loci that contribute to risk including several copy number variants (CNV) of large effect and hundreds of associated loci of small effect. However, in only a few cases has a specific gene been clearly identified. Rare CNV that affect only a single gene offer a potential avenue to discovering specific SCZ risk genes. Here, we use CNV generated from exome-sequencing of 4,913 SCZ cases and 6,188 controls in a homogenous Swedish cohort to assess the contribution of single-gene deletions and duplications to SCZ risk. As previously seen, we found an excess of rare deletions (p = 0.0004) and duplications (p = 0.0006) in SCZ cases compared to controls. When limiting to only single-gene CNV we identified nominally significant excess of deletions (p = 0.04) and duplications (p = 0.03). In an effort to increase the number of single-gene CNV, we reduced strict filtering criteria but required support from two independent CNV calling methods to create an expanded set that showed a significant burden of deletions in 11 out of 22 gene sets previously implicated in SCZ and in the combined set of genes across those sets (p = 0.008). Finally, for the significantly enriched set of voltage-gated calcium channels, we performed an extensive validation of all deletions generated from exome-sequencing as well as any deletion with evidence from previously analyzed genotyping arrays. In total, 4 exonic, single-gene deletions validated in cases and none in controls (p = 0.039), of which all were identified by exome-sequencing. Broadly, these results point to the potential contribution of single-gene CNV to SCZ and the added value of a deeper dive into CNV calls from exome-sequencing.

## Introduction

Schizophrenia (SCZ) is a highly heritable psychiatric disorder that causes substantial morbidity, mortality, and personal and societal costs(1–4). In order to increase our understanding of the biological basis of SCZ, it is important to identify genetic variation that influences risk. Copy number variants (CNV) are appealing as they directly alter gene dosage or structure and thus provide an interpretable effect on gene function. SCZ cases have been repeatedly shown to carry a greater burden of large/rare CNV (>100 kb and <1%), particularly deletions affecting brain-expressed genes. Multiple rare recurrent CNV with substantial effects on risk for SCZ (genotypic relative risks 4-20) have been identified (e.g., 16p11.2 and 22q11.21)(5–10). Most of these known CNV risk loci are megabase-sized and affect the dosages of many genes, but if specific genes contributing to risk could be identified it would aid our understanding of the neurobiology of the disorder. Thus far, only a few specific genes from genetic studies of CNV and SNV have been implicated in SCZ: *NRXN1*(11)*, TOP3B*(12), *RBM12*(13) and *SETD1A*(14), all of which provided novel insights into SCZ pathophysiology. Therefore, gene-focused CNV evaluation in large samples with high resolution capture is needed.

The majority of CNV have a small genomic footprint(15–18) and, due to either technological limitations or cost associated with identifying them, their contribution to SCZ remains unknown(19). Commercial microarrays are limited by probe density and are largely incapable of detecting CNV below 10 kb while also having low specificity for CNV between 10 and 100 kb(20). CNV detection from whole genome sequencing represents a substantial improvement, but remains expensive and is presently infeasible for large samples. Whole exome sequencing (WES) data can be used to identify CNV impacting exons(21). These data, while noisy from dependence solely on read depth and lacking exact breakpoints from the discrete nature of exons, can be used to increase power to identify smaller CNV affecting single genes that may be more interpretable in their contribution to SCZ risk.

Here, we have performed a comprehensive analysis of WES data from the Swedish Schizophrenia Study, which included 4,978 schizophrenia cases and 6,256 controls(22). Our goals were to evaluate the overall impact of single-gene CNV on SCZ risk, and to discover risk loci resulting from changes in copy number in specific genes that could lead to improved mechanistic understanding of SCZ. The Swedish sample is well suited for this study given its national sampling framework and relative homogeneity. All samples were also genotyped with GWAS genotyping arrays and Illumina exome arrays(23,24) providing additional data to follow up and validate CNV calls.

## Methods

### Sample description

We obtained DNA extracted from venous blood samples from 11,234 Swedish participants (mean age at time of sample collection, 55 years). This cohort included 6,256 controls and 4,978 SCZ cases. An additional 1,172 Swedish samples were included in generating and cleaning exome-sequencing CNV calls to improve estimates of copy number and frequency but were removed before analyses (total N: 12,384). All procedures were approved by ethical committees in Sweden and the US, and all subjects provided written informed consent. Genomic investigation of each subject was done using independent technologies including GWAS array genotyping(23), exome array genotyping(19), and exome sequencing(22,25). All genotyping and sequencing was conducted at the Broad Institute. Rare CNV data from GWAS genotyping array and exon genotyping arrays had been previously generated(19,24), and is briefly described in the supplementary material. Exome-sequencing based CNV were generated for this analysis and have not previously been reported on. Individuals already known to be carrying large CNV were included in all analyses. All genomic locations are given in NCBI build 37/UCSC hg19 coordinates.

### CNV calling and QC using XHMM

We ran XHMM (eXome-Hidden Markov Model) as previously described(21,26), including calculating mean per-base coverage across 189,894 targets (sequences designed for capture, predominantly exons) using GATK DepthOfCoverage. A total of 14,555 targets were excluded before CNV calling due to: mean sequencing depth <10x, low complexity sequence (as defined by RepeatMasker) in >25% of its span, GC content <10% or >90%, and spanning < 10 bp or >10 kb. The resulting sample-by-target read depth matrix was scaled by mean-centering the targets, after which principal component analysis (PCA) of the matrix was performed. To normalize the data, the top 109 principal components (those with variance >70% of the mean variance across all components) were removed from the data to account for systematic biases at the target- or sample-level, such as GC content or sequencing batch effects. Additional targets (n = 37) were removed if variance in read depth remained high after normalization (standard deviation >50). CNV were called using the Viterbi hidden Markov model (HMM) algorithm with default XHMM parameters, and XHMM CNV quality scores (SQ) were calculated using the forward-backward HMM algorithm. For any CNV detected in at least one individual we statistically genotyped all samples using the same XHMM quality scores and output as a single VCF file. Twenty-two samples failed CNV calling from XHMM due to low overall read depth, A total of 175,303 targets were used to call CNV across 12,384 samples after all filtering. CNV from sex chromosomes would be inaccurately called since males and females were run together and so were removed from analyses.

There were 494,403 autosomal CNV called by XHMM before any further filtering. We removed 115 individuals with > 3 standard deviations from the mean in total number of CNV (71.5) or total genomic content affected by CNV (6,529 kb). After sample outlier removal, 484,940 CNV (SQ > 0) were used to develop a frequency filter, and we retained only CNV seen in less than 1% of individuals (<0.5% minor allele frequency) across the entire sample as used in previous work on rare CNV. To account for the discrete nature of exons, each target was numbered sequentially regardless of genomic coordinates and frequency filtering was done using the sequential target information before mapping targets to genomic positions. After frequency filtering there were 51,812 CNV total with a mean of 4.3 per individual ranging from 1-107. After performing the recommended filter (SQ ≥ 60), 14,243 CNV remained (we refer to this dataset going forward as the “exome QC” dataset).

### Expanded single-gene CNV dataset integrating XHMM and ExomeDepth

In analyzing XHMM data, we observed that filtering on increasing quality scores (SQ) disproportionately removed shorter CNV. In an effort to quantify the proportion of shorter CNV with lower quality scores that were likely to be real we used exome-sequencing data from a set of 624 trios(27) and calculated transmission as a function of quality score and minimum number of targets required per CNV. These data were processed using the same software (XHMM) and QC as described above. We focused on rare CNV (< 0.1%) to avoid counting transmissions arbitrarily. At the recommended filtering thresholds (SQ ≥ 60, minimum 3 exons) we calculated a transmission rate of 0.42 (64 maternal CNV, 26 transmitted; 71 paternal CNV, 30 transmitted). When we looked at only those CNV having a single supporting exon and no minimum SQ (i.e. all called CNV) we saw a significant increase in CNV but an expected reduction in transmission rate of 0.114 (449 maternal CNV, 55 transmitted; 575 paternal transmitted, 62 transmitted). However, this transmission rate still suggests that potentially 20+% of these “low quality” events may be real. As an effort retain the true shorter events while removing as many of the false positive CNV calls as possible we required additional support from an independent approach ExomeDepth(28). Briefly, ExomeDepth selects a small set of individuals to be used as the reference group for CNV inference. These individuals are selected for having similar sequencing properties as the individual being called and this selection process is independently performed for each individual. We therefore called CNV within experimental plates of 96 individuals that were processed and sequenced at the same time. We generated CNV calls for all individuals using ExomeDepth. In total, we called CNV for 12,313 samples totaling 1,915,300 CNV with a mean of 155.5 per individual and ranging from 1-811. We retained any XHMM call with SQ ≥ 60 and any CNV called by both ExomeDepth and XHMM regardless of quality score (we refer to this set going forward as the “expanded exome” dataset). For comparison, 92% of the high-quality calls from the exome QC dataset were also called by ExomeDepth whereas only 20% of CNV affecting a single exon and low SQ (< 30) were called across both methods. In total, the expanded exome dataset had 24,843 CNV compared to 14,243 in the exome QC dataset. Using the union of the two approaches allows us to expand our set of shorter CNV while retaining only those with the most support.

### CNV burden and association analyses

We performed several burden and association analyses using Plink(29). For burden analyses, we performed empirical permutation (n = 10,000) of case/control label where permutation was performed within sequencing batch to account for any batch effects. For gene-based tests, we performed the same empirical permutation procedure defined above for burden tests. CNV were considered to affect a gene if there was any overlap of the genomic coordinates of the CNV and the gene. For gene-set tests, we used a regression framework built into Plink(30) that tests whether cases carry more CNV in the set of genes compared to all genes after covarying for number and amount of CNV.

### Incorporating CNV from previously run genotyping arrays of the same individuals

To maximize the sensitivity to detect gene/exon level CNV, we constructed a union CNV call set by combining the CNV data from GWAS array, exon array, and our expanded exome CNV dataset. We first created a database of all non-redundant CNV, where, for each CNV record, we indicated (1) how many platform(s) had identified the CNV; (2) which specific platform(s) had identified the CNV; (3) the coordinates of CNV from each platform. We considered two CNV redundant if they have the same direction of the copy number change and they overlapped more than 50% of their lengths. Details for this “exome plus array” dataset are described in supplemental materials (**Tables S3-S4**, **Figures S1-S2**).

### Validation of CNV

We attempted validation on 55 deletions from the genotyping and exome CNV dataset that affected any calcium channel gene (N = 26 genes) using a combination of both quantitative PCR (qPCR) and NanoString nCounter technology. First, qPCR was used to verify CNV detected in calcium channel genes *CACNA2D3*, *CACNA1B*, *CACNA2D4* and *CACNG2* (**Table S5**). Several predesigned TaqMan Copy Number Assays were run in quadruplicate along with the internal RNase P Copy Number Reference Assay according to manufacturer’s instructions (Applied Biosystems, Foster City, CA). Briefly, 20 µl reactions containing 1 µl DNA (5 ng), 10 µl of 2X Taqman Genotyping Master Mix, 1 µl of one target CNV assay and 1 µl of RNase P reference assay were mixed. All qPCR reactions were run on a Life Technologies StepOnePlus machine with the following thermal cycling conditions: 95°C for 10 min, followed by 40 cycles of 95°C for 15 s and 60°C for 1 min. Samples included all suspected CNV carriers for each gene, regardless of case or control status, as well as four presumed two-copy controls per gene.

Second, for a larger scale validation of deletions in 48 additional samples, we used Nanostring nCounter technology. For each CNV, two probes were designed and analyses were performed according to manufacturer instructions. In brief, a spike-in plasmid of known amount was used to control for variability in DNA quantity across all samples and additional controls ensured optimal hybridization and purification efficiency. After hybridization and removal of excess probes, the probe/target complexes were aligned and immobilized in the nCounter Cartridge, and imaged in the nCounter Digital Analyzer for detection of CNV. In a previous study, we examined nCounter’s CNV calling accuracy by testing 37 known CNV in 384 samples and found 97% concordance in CNV calls.

## Results

### Exome-sequencing CNV demonstrate high concordance with genotyping array based CNV while contributing substantial numbers of novel variants

We generated CNV calls using XHMM for 4,913 SCZ cases and 6,188 controls resulting in a total of 14,243 rare (present in less than 1% of individuals) and high quality (SQ ≥ 60) CNV (“exome QC dataset”). In a comparison to previously published work on this sample where CNV were generated from genotyping arrays(24) (see Supplementary Methods) we identified 78% of the array-based CNV in the exome QC dataset. More interestingly, 75% of the exome QC calls were not seen in the array-based call set at all. Individuals carried, on average, 2.2 times more CNV in the exome QC dataset than in the array-based call set (1.28 versus 0.59 CNV). This comparison is described in more detail(21). Specific to this work, 53% of exome QC CNV overlapped a single protein coding gene and, of those, only 12.6% were included in the previous work on this sample leaving 87.4% or 6,622 single-gene CNV to be analyzed for the first time here.

### Significant burden of exome-sequencing based CNV in SCZ including among single-gene CNV

We first assessed the burden of all CNV in the exome QC dataset to SCZ. Utilizing empirical permutation of case/control label (see Methods) we identified a significant increase in the numbers of deletions (case rate: 0.56, control rate: 0.51, p = 0.0004) and duplications (case rate: 0.78, control rate: 0.72, p = 0.0006) in SCZ cases compared to controls as seen previously in this sample(24). To identify the contribution of the novel CNV in our exome QC dataset, we performed the same burden test using only CNV new to this analysis and not previously called by arrays in previous work. Here, we again saw significant burden in cases for both deletions (case rate: 0.48, control rate: 0.45, p = 0.0114) and duplications (case rate: 0.56, control rate: 0.51, p = 0.0003). The exome QC CNV are substantially shorter and therefore more likely to affect only a single gene. We tested whether burden of CNV was primarily driven by larger events affecting multiple genes or if single-gene CNV were contributing. We identified a significant but modest burden specific to single-gene deletions (case rate: 0.36, control rate: 0.34, p = 0.0395) and duplications (case rate: 0.34, control rate: 0.32, p = 0.0332) in SCZ cases compared to controls which was not as significant as the larger events (***Figure 1, Table 1***).

**Table 1.** CNV burden results stratified by CNV type (deletions, duplications), number of genes affected (all, single gene or multiple genes) and whether the CNV was unique to our exome-sequencing call set or was identified in previous array-based CNV work. Bolded p-values are less than 0.05.

|  |  | *Deletions* |  |  |  | *Duplications* |  |  |  |
| --- | --- | --- | --- | --- | --- | --- | --- | --- | --- |
|  |  | N | Case rate | Control rate | P | N | Case rate | Control rate | P |
| All | All | 5900 | 0.56 | 0.51 | **0.0004** | 8343 | 0.78 | 0.73 | **0.0006** |
|  | New | 5101 | 0.48 | 0.45 | **0.0114** | 5925 | 0.56 | 0.51 | **0.0003** |
|  | Previously called by genotyping arrays | 799 | 0.08 | 0.06 | **0.0002** | 2418 | 0.23 | 0.21 | 0.1396 |
| Single gene | All | 3894 | 0.36 | 0.34 | **0.0395** | 3680 | 0.34 | 0.32 | **0.0332** |
|  | New | 3530 | 0.33 | 0.31 | 0.0543 | 3092 | 0.29 | 0.27 | **0.0162** |
|  | Previously called by genotyping arrays | 364 | 0.03 | 0.03 | 0.1998 | 588 | 0.05 | 0.05 | 0.6180 |
| Multiple genes | All | 1773 | 0.18 | 0.14 | **0.0001** | 4516 | 0.42 | 0.39 | **0.0030** |
|  | New | 1339 | 0.13 | 0.11 | **0.0174** | 2695 | 0.25 | 0.23 | **0.0076** |
|  | Previously called by genotyping arrays | 434 | 0.05 | 0.03 | **0.0001** | 1821 | 0.17 | 0.16 | 0.0756 |

**Figure 1.**
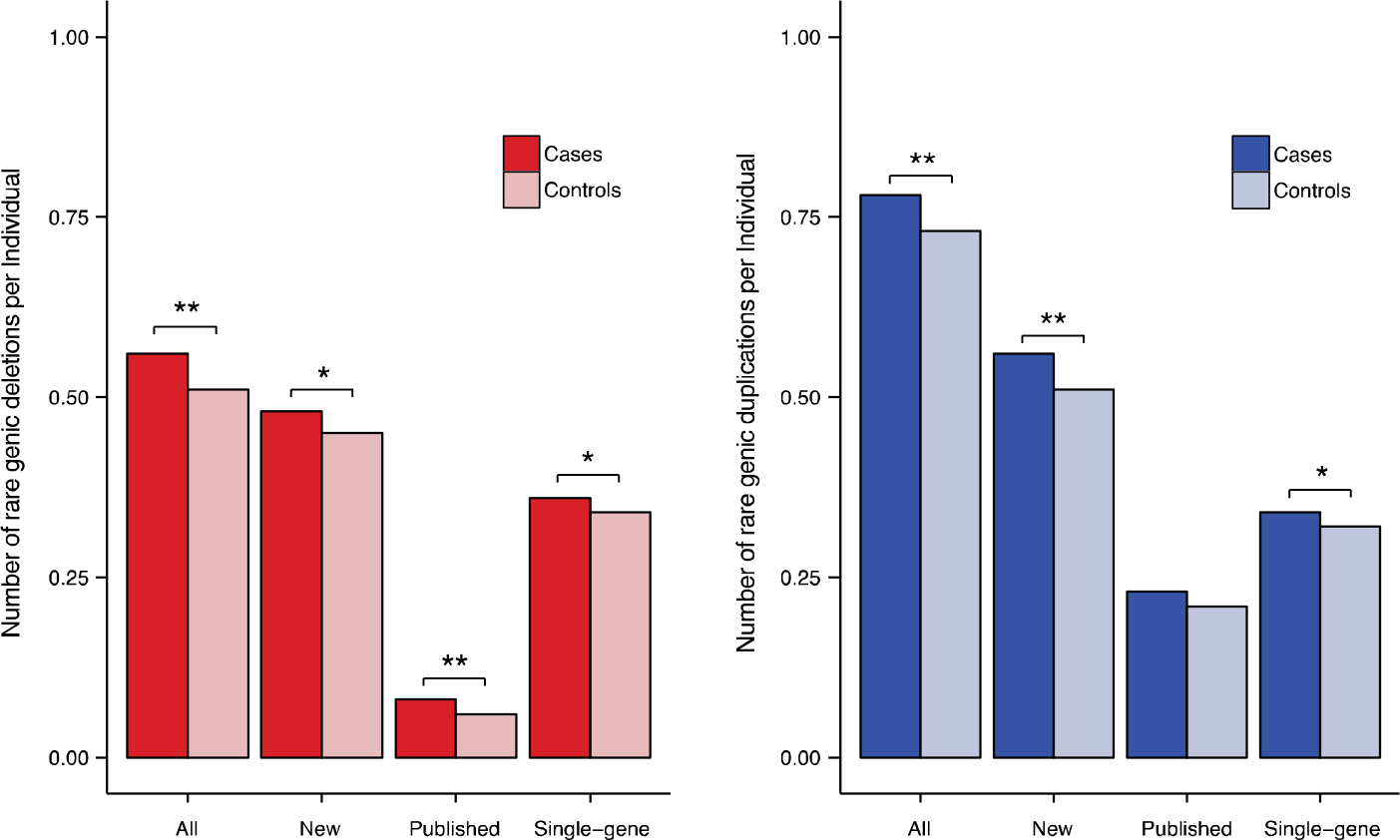
Burden tests across all high confident exome-seq CNV calls (all), those not previously analyzed from genotyped arrays (new), those previously published (published) and only those CNV affecting a single protein coding gene (single-gene). Deletions are in red (left) and duplications are in blue (right). Significance is represented as p < 0.05 (*), p < 0.001 (**).

### Expanding the set of potential single-gene CNV and testing for excess in specific genes

We next sought to test whether CNV could implicate specific genes using both the exome QC dataset as well as an expanded exome dataset created to increase the proportion of shorter CNV which our QC filters were disproportionately removing (see Methods). Briefly, we created this dataset by integrating CNV calls from both XHMM and ExomeDepth(28) retaining CNV if detected by both methods regardless of XHMM quality scores or if detected only by XHMM at our previous filtering threshold (SQ ≥ 60). In total, our “expanded exome dataset” includes an additional 10,600 CNV (total: 24,843) substantially increasing the proportion of shorter events. Individual genes were tested for excess of deletions or duplications using empirical permutation. After 10,000 permutations in our exome QC dataset, 21 genes were significantly enriched for duplications and 40 genes were significantly enriched for deletions in cases compared to controls after multiple test correction. However, all significant genes fell into two regions driven by already known large SCZ risk CNV, 16p11.2 (duplications) and 22q11.2 (deletions) leaving no novel genes identified (***Figure 2***). Finally, using our expanded exome dataset we again tested for enrichment of deletions and duplications in specific genes. No gene was significant after correction for multiple testing with the most significant genes again being driven by the larger 16p11.2 or 22q11.2 CNV.

**Figure 2.**
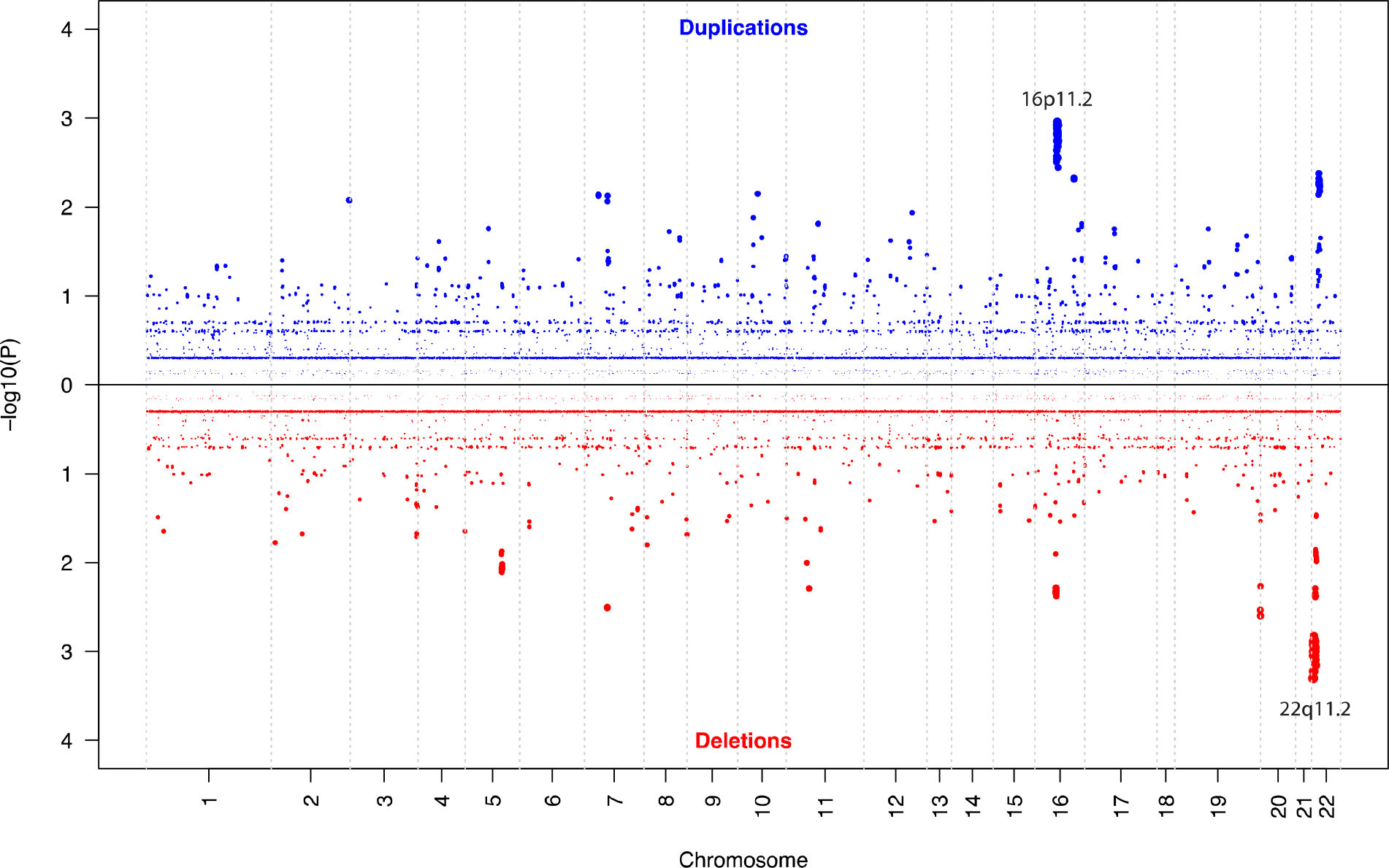
Gene-based Manhattan plot of duplications in blue (top) and deletions in red (bottom). Genes in most significant regions are labeled by known CNV in that region.

### Testing contribution of only single-gene CNV to previously implicated SCZ gene sets

In the absence of any single novel gene being significant we sought to implicate particular biological pathways from those CNV affecting only single genes. Here, we again leveraged our expanded exome dataset but filtered out any CNV that affected more than one gene. In total, there were 14,091 CNV affecting only a single protein-coding gene (7,423 deletions, 6,668 duplications), all larger CNV including those previously implicated were excluded from these analyses. We tested genesets previously implicated in SCZ to address whether single-gene CNV are contributing a significant proportion of risk, as opposed to being predominantly driven by larger multi-gene CNV. The sets tested included genes previously implicated directly in SCZ (GWAS loci(31), *de novo* variants(32), CNV regions(7)), synaptic function(33) (ARC, mGluR5, NMDAR, PSD95), calcium channels(25) (CAV2, Voltage-gated), secondary sets (FMRP targets(22,25), ASD/DD/ID *de novo*(32), essential genes(34), constrained genes(35), RBFOX related genes(22) and antipsychotic targets(36)). All sets combined included 8,970 genes and showed significant excess in cases for deletions (p = 0.008) but not duplications (p = 0.186). Among each gene set tested individually, none surpassed a Bonferroni corrected p-value of 0.001 for the 44 tests performed. However, we identified nominally significant enrichment of single-gene deletions in half (11 out of 22) of the sets (***Table 2***).

**Table 2.** Geneset CNV results for single-gene CNV in expanded dataset. Bold represents path ways with p-value < 0.05.

| Group | Set | N genes | *Single gene CNV* |  |
| --- | --- | --- | --- | --- |
|  |  |  | P del | P dup |
| SCZ sets | PGC2 SCZ 108 loci | 329 | 0.4199 | 0.7021 |
|  | SCZ de novo LoF | 87 | **0.0296** | 0.09109 |
|  | SCZ de novo NS | 611 | 0.2761 | 0.9024 |
|  | PGC2 16 CNV | 175 | **0.005499** | 0.9989 |
|  | PGC2 16 CNV (deletions) | 78 | 0.2594 | 0.9289 |
|  | PGC2 16 CNV (duplications) | 111 | **0.007699** | 0.579 |
| Synaptic Sets | ARC | 28 | 1 | 0.3543 |
|  | mGluR5 | 39 | 0.2425 | 0.8117 |
|  | NMDAR_network | 61 | **0.0335** | 0.8099 |
|  | PSD-95_(core) | 65 | 0.06969 | 0.8984 |
| Calcium Channel Sets | CAV2 | 206 | **0.0347** | 0.4065 |
|  | CAV2 Ion | 44 | 0.1738 | 0.2395 |
|  | Voltage-gated_Calcium_Channel_Genes | 26 | **0.008199** | 0.2792 |
| Secondary Sets | FMRP-targets | 788 | **0.0205** | 0.5018 |
|  | ASD de novo | 1080 | 0.05209 | 0.4664 |
|  | DD de novo | 1271 | **0.0161** | 0.966 |
|  | ID de novo | 350 | 0.08979 | 0.5774 |
|  | Antipsychotic targets | 347 | **0.0268** | 0.9338 |
|  | Essential genes | 3915 | 0.06219 | 0.1975 |
|  | Rbfox | 2737 | **0.005799** | 0.2056 |
|  | LoF intolerant (pLI > 0.9) | 3488 | **0.0163** | 0.243 |

### Broad scale exploration of CNV in calcium channel genes combining both the expanded exome dataset and array-based calls

Of the most significant gene sets, we selected the voltage-gated calcium channel set for a full-scale validation since it represented an approachable number of genes to validate all overlapping CNV comprehensively and had significant prior supporting literature. Across the 26 genes, we identified 6 deletions in cases and 0 in controls from our expanded exome dataset (***Figure 3***). Since validation with an independent technology is considered the gold standard for CNV work, we attempted to validate these deletions using quantitative PCR (qPCR). Four of the deletions validated, two identical single exon deletions in *CACNA2D3* did not validate (these two did not surpass filtering thresholds to be included in the exome QC dataset). Since we had additional CNV data from genotyping arrays, we wanted to validate a larger set of calcium channel deletions to more comprehensively catalog the contribution of deletions in these genes to risk of SCZ in this sample. We identified a comprehensive set of deletions across our exome plus array dataset (see Methods and Supplementary Methods) overlapping any voltage-gated calcium gene. In total, we identified 55 deletion calls in 55 different samples which we validated using NanoString nCounter technology (see Methods). Of these, 34 were located over three common, intronic copy number polymorphisms (all of which validated). Of the 21-remaining rare-variant calls, 6 validated (see ***Table 3***). The low validation rate is representative of our decision to take all CNV calls with limited evidence and not filter on confidence. Nearly all of CNV that did not validate were low quality calls from the genotyping arrays. Of the 6 validated deletions, 4 were single-gene and directly affected exons and all were seen in cases. All 4 were identified in the expanded exome dataset. We further validated a non-exonic deletion and a multi-gene deletion in controls identified from genotyping arrays. After validation, we were left with 4 single-gene deletions in cases and 0 in controls (p = 0.039).

**Table 3.** List of all rare deletions overlapping the 26 voltage-gated calcium channel genes that validated including one that did not overlap an exon and one that was not single-gene. Four of the 6 single-gene deletions identified in the geneset analyses and in figure 3 validated (CACNA2D3 did not).

| Status | Chr | Start | Stop | Size (bp) | Gene | Single-gene? | Exonic? |
| --- | --- | --- | --- | --- | --- | --- | --- |
| case | 9 | 140866027 | 141004005 | 137978 | CACNA1B | yes | yes |
| case | 9 | 140846726 | 141016451 | 169725 | CACNA1B | yes | yes |
| case | 12 | 1949932 | 1965357 | 15425 | CACNA2D4 | yes | yes |
| case | 22 | 36960396 | 36960935 | 539 | CACNG2 | yes | yes |
| control | 3 | 54262746 | 54316431 | 53685 | CACNA2D3 | yes | **no** |
| control | 7 | 79818265 | 82072777 | 2254512 | CACNA2D1 | **no** | yes |

**Figure 3.**
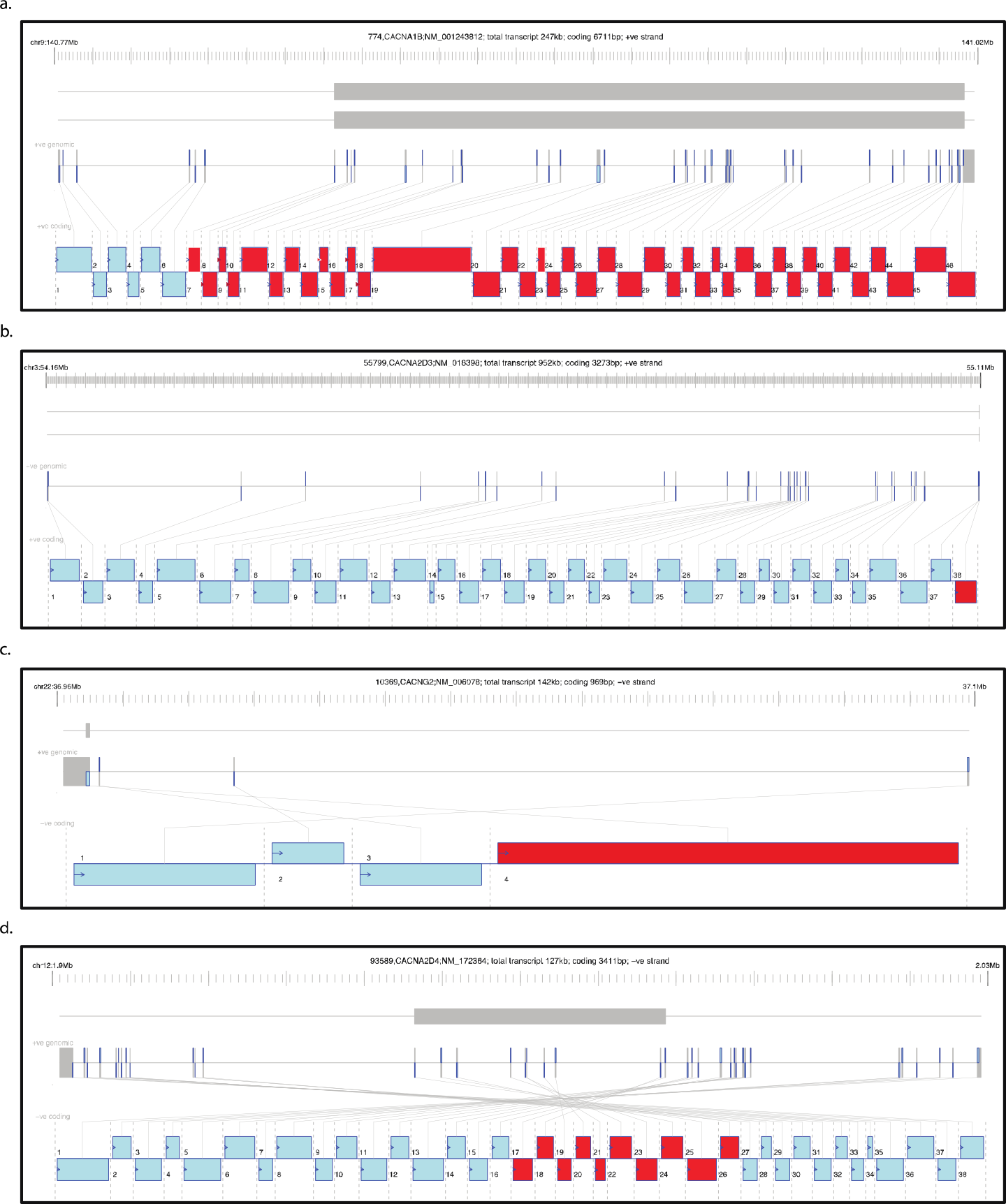
Gene model plots for each of the 4 genes and 6 deletions identified in voltage-gated calcium channel genes. Upper grey bars portray deletion in genomic space, below that is the gene model in genomic space. The bottom bars represent the exons as transcribed, red indicates exons that were deleted. All deletions replicated except the two shown in panel b.

## Discussion

This study represents an evaluation of smaller CNV in a large SCZ sample. We found that, independent of larger events, deletions of single genes may contribute to schizophrenia risk through a number of biological pathways previously identified for SCZ. In particular, we identify and validate a small number of deletions in voltage-gated calcium channels that are enriched in SCZ cases compared to controls. We also demonstrate the utility of exome-sequencing to identify shorter, single-gene CNV and the potential to improve the resolution of those events through combining multiple methods for further study.

The majority of contribution of CNV to SCZ to date has been in the form of large (>100kb) and rare CNV both in specific loci or across the genome(7). The ability to determine the contribution from shorter CNV has been both technologically limited by the use of genotyping arrays but also biologically up for debate as few single genes have been implicated in SCZ risk and nearly all risk increasing CNV affect many genes. Here, we point to the potential contribution of single-gene CNV to risk for SCZ. This contribution can be identified both genome-wide but also within genes having been previously implicated from other studies of genetic variation including synaptic genes, genes having *de novo* mutations in SCZ, DD, ASD or ID, conserved genes and gene targets of antipsychotics. This work points to a confluence of evidence that these gene sets are relevant for schizophrenia biology. We did not identify any specific gene that was significantly associated after correction for multiple testing. Given other studies of rare variation in complex diseases with similar sample sizes, this is not surprising(24) but does suggest that combining CNV data with SNV data could improve power to implicate specific genes and robust approaches to combine these classes of variation are needed.

Calcium channel genes have been implicated in psychiatric disease risk, including in SCZ for many years. Studies to date from the genetics of SCZ have implicated particular loci and the geneset as a whole. Here, we show an excess of single-gene CNV in calcium channels that remains after qPCR validation. Given the importance of this gene set and the relative size, we also performed a larger validation of deletions using a higher throughput method confirming the 4 qPCR validated single-gene deletions in cases using a different method as well as validating several common CNV, one >2Mb deletion in a control and one deletion that did not overlap an exon in a control. Our results suggest that deleting a single calcium channel gene may be relevant for SCZ risk however substantially more data will be required to confirm this finding.

We show that exome-sequencing can provide a substantial number of novel CNV that are not captured by genotyping arrays and are predominantly affecting only a single gene. Further, this work points to the existence of likely many real single-gene CNV that are filtered out by default filtering criteria and only by combining multiple currently existing approaches can we capture a reasonable set of the true calls without including too much noise. While exome-sequencing can substantially improve resolution of CNV calling it is not without its weaknesses and limitations that become even clearer as CNV get smaller. Whole-genome sequencing will offer the best resolution to confidently identify single-gene CNV but is still prohibitively expensive for most labs and hundreds of thousands of exome sequences currently exist, and many more are being generated, making CNV calling from exome-sequencing still important. We believe there are opportunities to improve the ability to call shorter CNV from exome-sequencing that are more sophisticated than merging call sets from multiple approaches.

Here, we demonstrate a potential role for single-gene deletions to contribute to SCZ risk through similar pathways as previously implicated. We perform a comprehensive validation of deletions in voltage-gated calcium channel genes and show an enrichment of these deletions in SCZ cases compared to controls. Finally, we demonstrate further utility for CNV generated from exome-sequencing and the ability to improve resolution of shorter events which could improve our ability to identify biological causes of diseases like SCZ.

## Acknowledgements

This work was supported by NIMH R21 MH104831 (JPS, JJC). PFS gratefully acknowledges support from the Swedish Research Council (Vetenskapsrådet, award D0886501). The Sweden Schizophrenia Study was supported by NIMH R01 MH077139.

