## Supplementary Information for "Characterization of single gene copy number variants in schizophrenia"

**Supplementary material**

### CNV data from GWAS genotyping array

#### Genotyping

As shown in ***Table S1***, samples were genotyped in six batches at the Broad Institute using Affymetrix 5.0 (3.9%), Affymetrix 6.0 (38.6%), and Illumina OmniExpress (57.4%) arrays according to the manufacturers’ protocols. Genotype calling and quality control was done in four sets corresponding to data from Affymetrix 5.0 (Sw1), Affymetrix 6.0 (Sw2-4), and the OmniExpress batches (Sw5, Sw6). SNP genotypes were called using Birdsuite (Affymetrix) or BeadStudio (Illumina).

#### Subject QC

SNP-based subject QC. A multi-step quality control (QC) procedure was carried out. The exclusionary measures of basic QC were: SNP missingness ≥ 0.05 (before sample removal); subject missingness ≥ 0.02; autosomal heterozygosity deviation; SNP missingness ≥ 0.02 (after sample removal); difference in SNP missingness between cases and controls ≥ 0.02; and deviation from Hardy-Weinberg equilibrium (*P* < 10^−6^ in controls or *P* < 10^−10^ in cases). After basic QC, we identified 39,239 SNPs suitable for robust relatedness testing (MAF >0.05, *r*^2^ < 0.05). Relatedness testing was done with PLINK ^7^ and pairs of subjects with $\hat{\pi}$ > 0.2 were identified and one member of each relative pair removed at random. Following QC, a total of 11,244 subjects remained and were used for subsequent CNV calling and analysis.

Intensity-based subject QC. Next we removed problematic arrays with high MAPD or high GC waviness using empirically derived thresholds (i.e., exceeding 3 standard deviations from the sample mean per array type). (MAPD is a measure of probe variance and is defined as the median of the absolute values of all pairwise differences between log_2_ ratios for a given genotyping array. GC waviness describes a spatial “wave” pattern in log_2_ ratios and is a systematic technical artifact observed in various array platforms.) Furthermore, we visualized pseudo-color images of all excluded arrays and a random sample of the good performing arrays to ensure the intensity-based QC worked properly.

CNV-load-based subject QC. Finally, we removed individuals who were outliers with respect to the total number or length of CNVs (>40 CNVs or total CNV spanning >6Mb). Thresholds were empirically derived as mean + 3xSD in the post-QC sample and by observing the distributions of these metrics across the entire dataset.

***Table S1.*** *GWAS array subject QC*

| **Feature** | **Sw1** | **Sw2** | **Sw3** | **Sw4** | **Sw5** | **Sw6** | **Total** |
| --- | --- | --- | --- | --- | --- | --- | --- |
| GWAS chip | affy5 | affy6 | affy6 | affy6 | ioexp | ioexp |  |
| Subjects (pre-QC) | 464 | 694 | 1,498 | 2,388 | 4,461 | 2,345 | 11,850 |
| After SNP-based QC | 427 | 643 | 1,356 | 2,262 | 4,361 | 2,195 | 11,244 |
| After intensity-based QC | 413 | 620 | 1,312 | 2,062 | 4,132 | 2,111 | 10,650 |
| After CNV-load-based QC | 413 | 616 | 1,310 | 2,058 | 4,130 | 2,109 | 10,636 |

#### CNV calling and QC

We used Birdseye to detect CNVs. We annealed adjoining CNVs that appeared to be artificially split by Birdseye by recursively joining CNVs if the called region is ≥ 80% of the entire region to be joined. CNV QC included removal of low-confidence CNVs with confidence scores < 10, spanning < 10 probes, or < 20 kb in length followed by removal of CNVs with > 50% reciprocal overlap with large genomic gaps (e.g., centromeres) or regions subject to rearrangement in white blood cells. We then imposed a 0.01 frequency threshold to extract rare CNVs. The final CNV data included a total of 5,997 deletions and 3,727 duplications in 4,719 cases with SCZ and 5,917 controls.

### CNV data from exon genotyping array

#### Genotyping

The same DNA samples were genotyped at the Broad Institute using the Illumina Infinium HumanExome BeadChip v1.0. Samples were placed on 96-well plates for processing and were processed within a relatively short time window. Majority of SNP genotypes were called using GenomeStudio v2010.3 with the calling algorithm/genotyping module version 1.8.4 using the custom cluster file StanCtrExChp_CEPH.egt, subsequent processing of genotype calling was done by zCall.

#### Subject QC

Subject QC was similar to GWAS arrays, which is based on SNP-based QC, intensity-based QC, and CNV-load-based QC. First, the 11,244 subjects that passed SNP-based QC filters as described in the previous section “GWAS array genotyping” were extracted. Next, low quality samples were excluded if they had extreme values for probe variance (i.e. LRR_standard deviation > 0.2, 95^th^ percentile or BAF_drift > 0.01, 95^th^ percentile), or were outliers with respect to the total number of CNV calls (>152, 95^th^ percentile).

***Table S2.*** *Exome genotyping array subject QC*

| **Feature** | **Sw1** | **Sw2** | **Sw3** | **Sw4** | **Sw5** | **Sw6** | **Total** |
| --- | --- | --- | --- | --- | --- | --- | --- |
| Subjects (pre-QC) | 464 | 694 | 1,498 | 2,388 | 4,461 | 2,345 | 11,850 |
| After SNP-based QC | 427 | 643 | 1,356 | 2,262 | 4,361 | 2,195 | 11,244 |
| After intensity- and CNV-load-based QC | 341 | 526 | 1,238 | 2,050 | 4,110 | 1,966 | 10,231 |

#### CNV calling and QC

CNV calling began with raw intensity data processing. A custom cluster file was created using the GenCall algorithm based on all samples. Normalized intensity values were obtained using Illumina’s GenomeStudio (v2010.3) with the calling algorithm/genotyping module (v1.8.4). For CNV calling, PennCNV (June 2011 version) was applied to the log R ratios (LRR) and B allele frequencies (BAF) calculated from the normalized intensity values. The default waviness correction and customized PennCNV parameters were used. CNV QC included removal of low-confidence CNVs with confidence scores < 10 or spanning < 10 probes, followed by removal of CNVs with > 50% reciprocal overlap with large genomic gaps (e.g., centromeres) or regions subject to rearrangement in white blood cells. We then imposed a 0.01 frequency threshold to extract rare CNVs. The final CNV data included a total of 4,707 deletions and 21,888 duplications in 3,962 cases with SCZ and 5,138 controls.

### Combining CNVs from multiple complementary platforms

In order to maximize the sensitivity to detect gene/exon level CNVs, we combined the post-QC CNV data from GWAS array, exon array, and exome sequencing and constructed a union callset of all non-redundant CNVs. The rationale for combing CNVs from multiple complementary platforms is illustrated in ***Figure S2***. We considered two CNVs redundant if they have the same direction of the copy number change and overlapped more than 50% of their length. For each CNV record, we indicated (1) how many platform(s) had identified the CNV; (2) which specific platform(s) had identified the CNV; (3) the coordinates of CNVs from each platform.

#### Subjects in the combined data

***Tables S3-S4*** and ***Figure S1*** show subject distribution with respect to the number of platforms (GWAS array, Exon array, or ExSeq) in the combined data.

***Table S3.*** *Subject distribution*

| **Feature** | **Sw1** | **Sw2** | **Sw3** | **Sw4** | **Sw5** | **Sw6** | **Total** |
| --- | --- | --- | --- | --- | --- | --- | --- |
| # Subjects with 1 platform | 14 | 49 | 25 | 39 | 27 | 25 | 179 |
| # Subjects with 2 platforms | 89 | 93 | 192 | 358 | 451 | 378 | 1,561 |
| # Subjects with 3 platforms | 322 | 494 | 1,139 | 1,859 | 3,880 | 1,786 | 9,480 |

***Table S4.*** *Subject distribution by case/control status*

|  | **Cases** | **Controls** | **Total** |
| --- | --- | --- | --- |
| Intersection | 4,188 | 5,292 | 9,480 |
| Union | 4,983 | 6,237 | 11,220 |

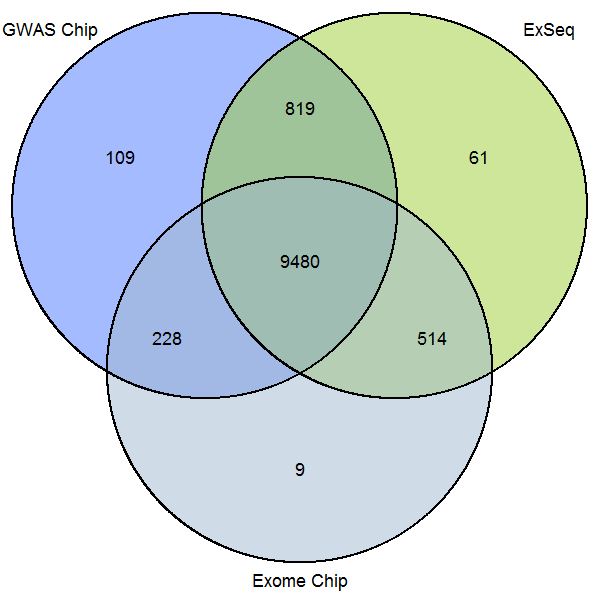

***Figure S1 Subject distribution across 3 platforms***

#### Illustration of the rationale

***Figure S2*** provides an example of a candidate single-gene CNV detected from the combined data but not using a single technology. VPS13B has deletions in 9 SCZ cases and 0 controls across all three platforms, and all verified by Q-PCR. VPS13B exemplifies the rationale behind combining CNVs from multiple complementary platforms to improve the sensitivity of CNV detection. This result did not replicate in 6,882 SCZ cases and 7,979 controls from 4 UK cohorts where we identified 2 SCZ deletions and 3 control deletions.

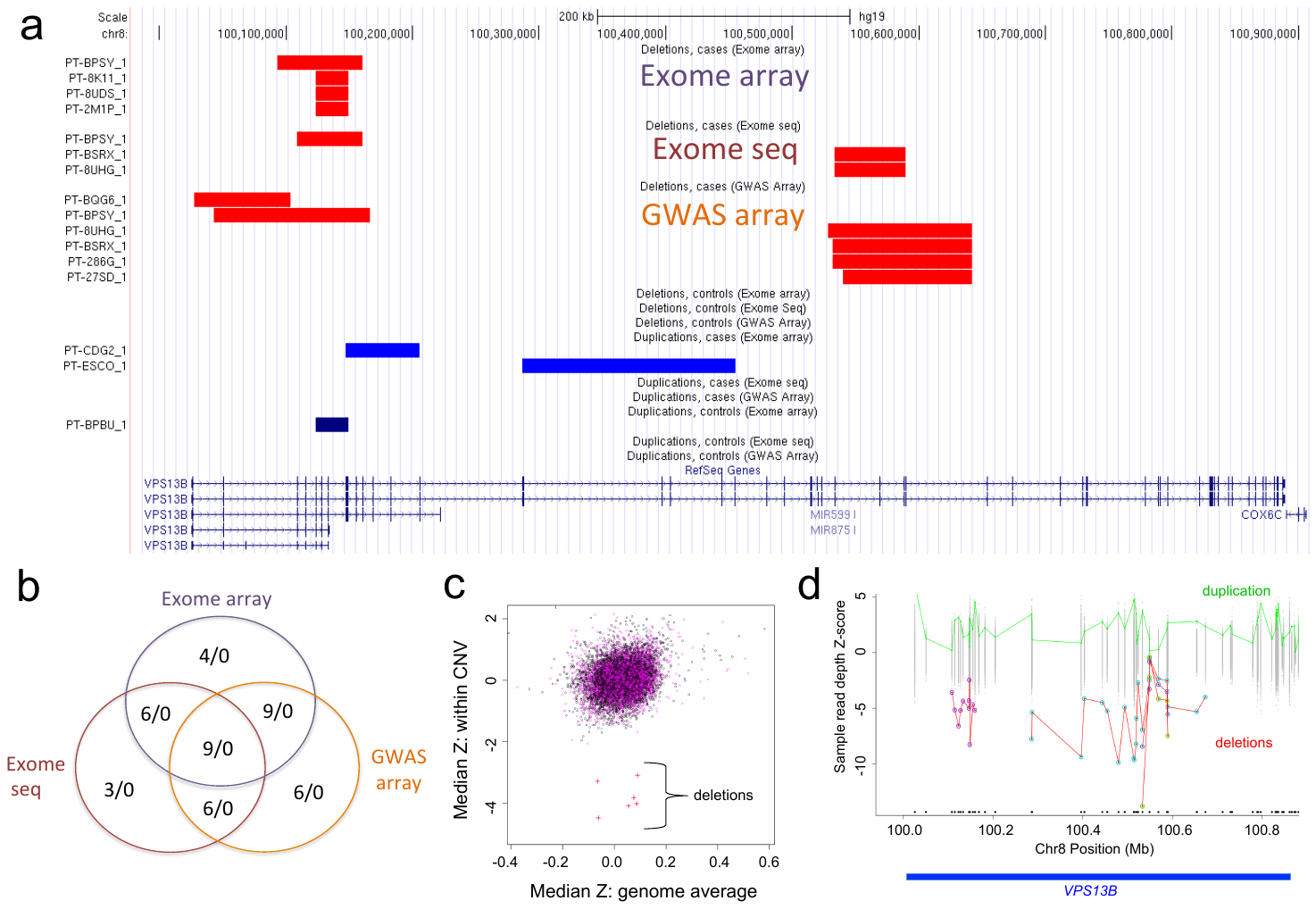

***Figure S2)*** VPS13B is a candidate single-gene CNV detected only from the combined CNV data. a) VPS13B has deletions in 9 SCZ cases and 0 controls across all methods. b) Within each technology, the case/control deletion results are suggestive; across technologies, they are compelling. c) Median Z score example (GWAS array). d) XHMM exome sequencing Z-score for VPS13B.

### Q-PCR copy number assays

| **Table S5.** Q-PCR copy number assays | | | |
| --- | --- | --- | --- |
| TaqMan assay ID | Gene | Chr | Position (hg19) |
| Hs02095527_cn | CACNA2D3 | 3 | 55108215 |
| Hs02095527_cn | CACNA2D3 | 3 | 55108215 |
| Hs02488682_cn | CACNA2D3 | 3 | 54872658 |
| Hs02095527_cn | CACNA2D3 | 3 | 55108215 |
| Hs03708188_cn | CACNA1B | 9 | 140940544 |
| Hs03708188_cn | CACNA1B | 9 | 140940544 |
| Hs01735338_cn | CACNA1B | 9 | 140777209 |
| Hs01987954_cn | CACNA1B | 9 | 140881248 |
| Hs02123786_cn | CACNA2D4 | 12 | 1965193 |
| Hs02123786_cn | CACNA2D4 | 12 | 1965193 |
| Hs01077756_cn | CACNG2 | 22 | 36960758 |
